## Supplementary figures and images for "Impairment of lipid homeostasis causes lysosomal accumulation of endogenous protein aggregates through ESCRT disruption"

### CRISPRi_LysoIP_UreaSDS_Silverstain_raw.tif

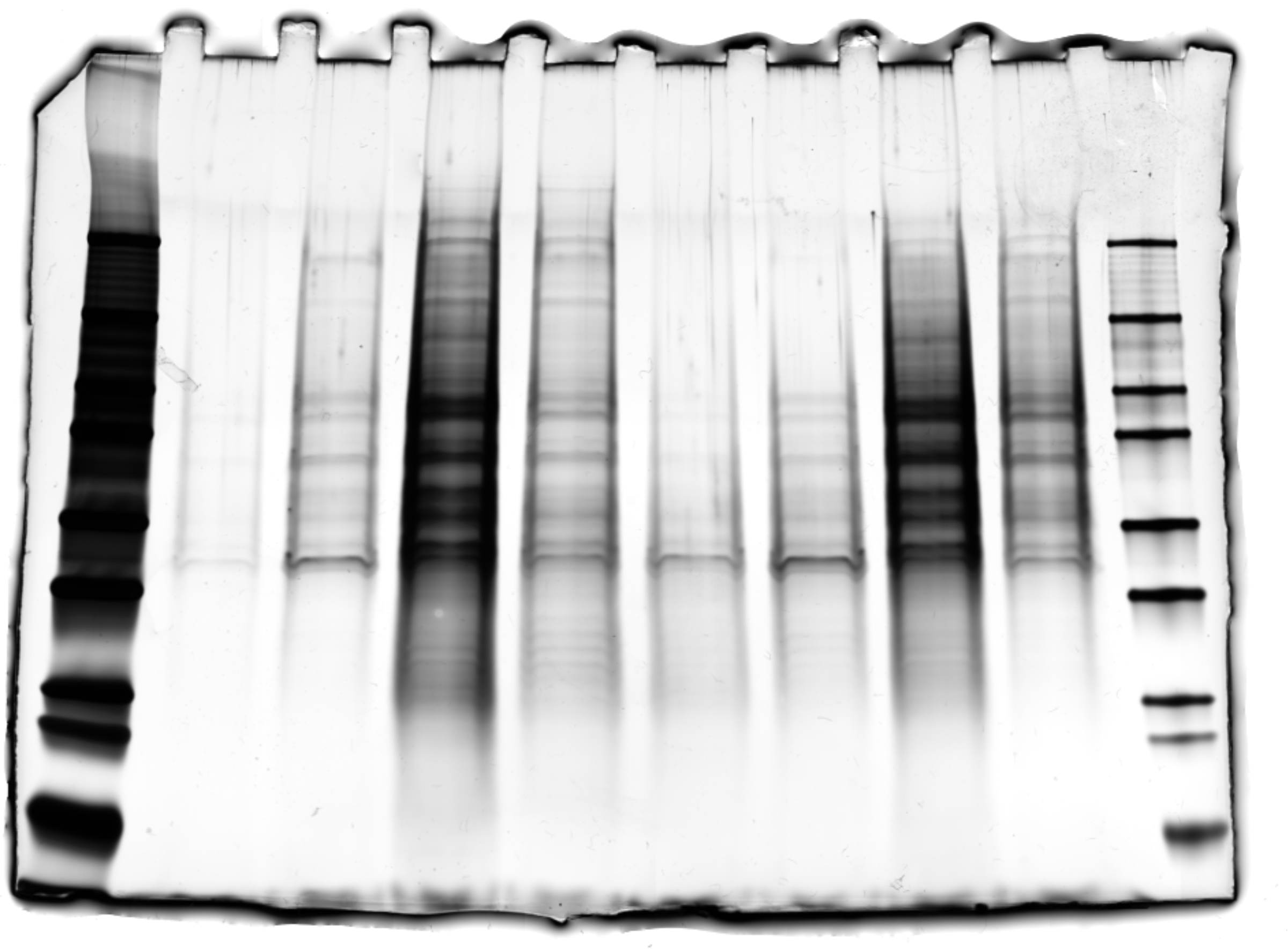

### Figure 3 - source data 1

#### Figure 3 – source data 2

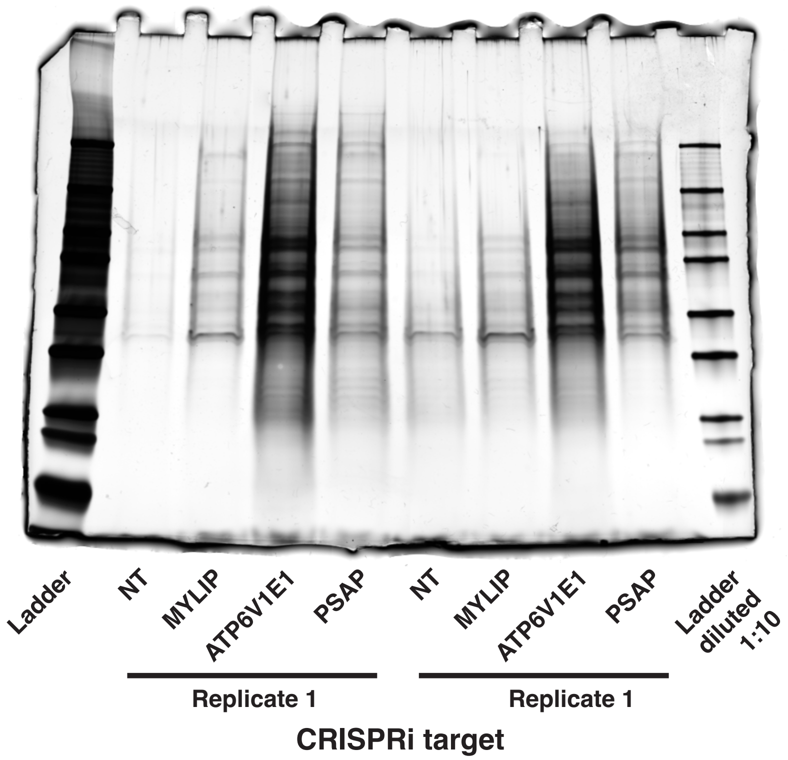


Ladder = BioRad Precision Plus Dual Color Protein Standards

### MG132_Bortezomib_Me4BoVS.tif

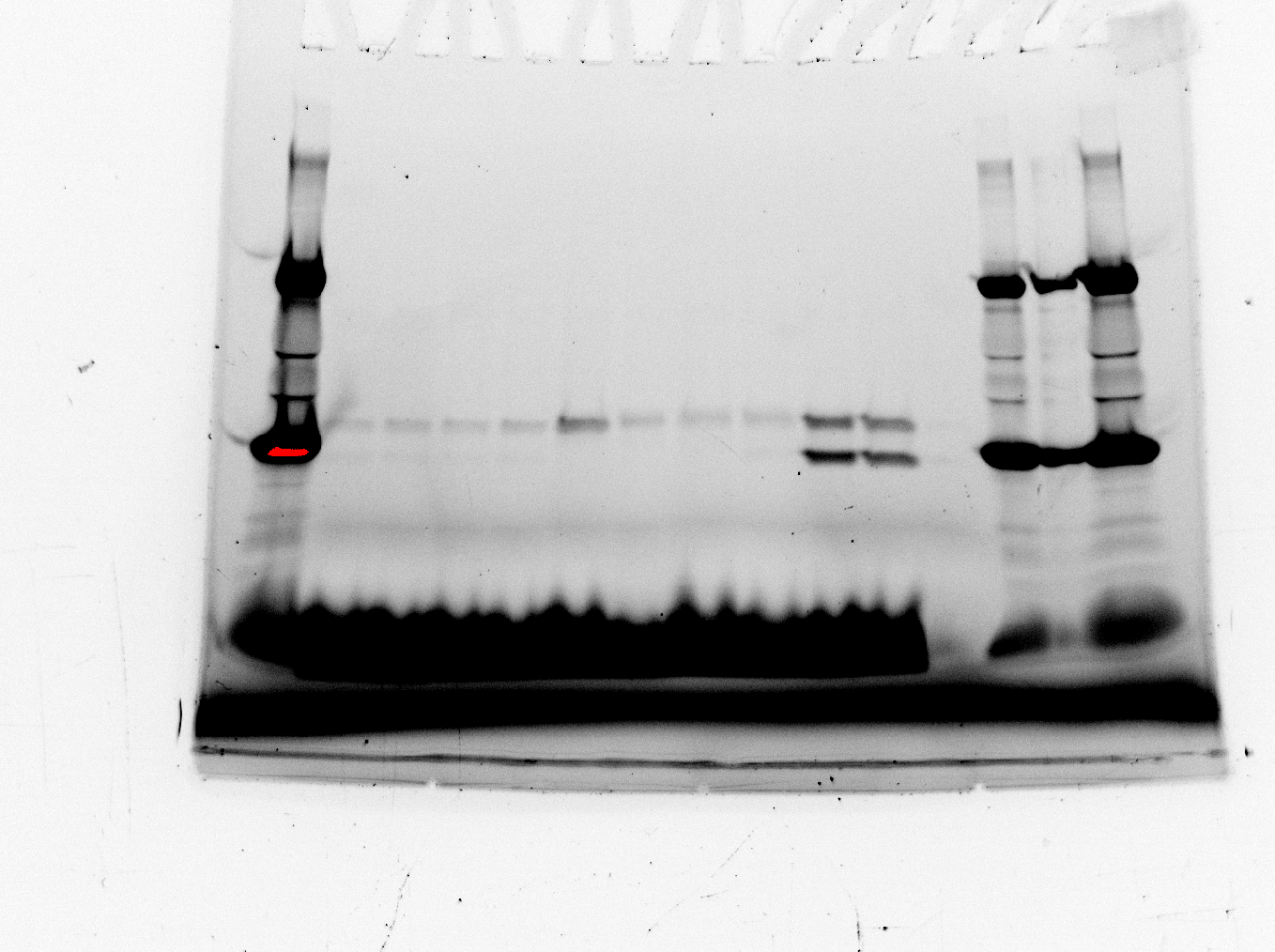

### MG132_series_Mock_and_Lyso-IP_Silverstain_raw.tif

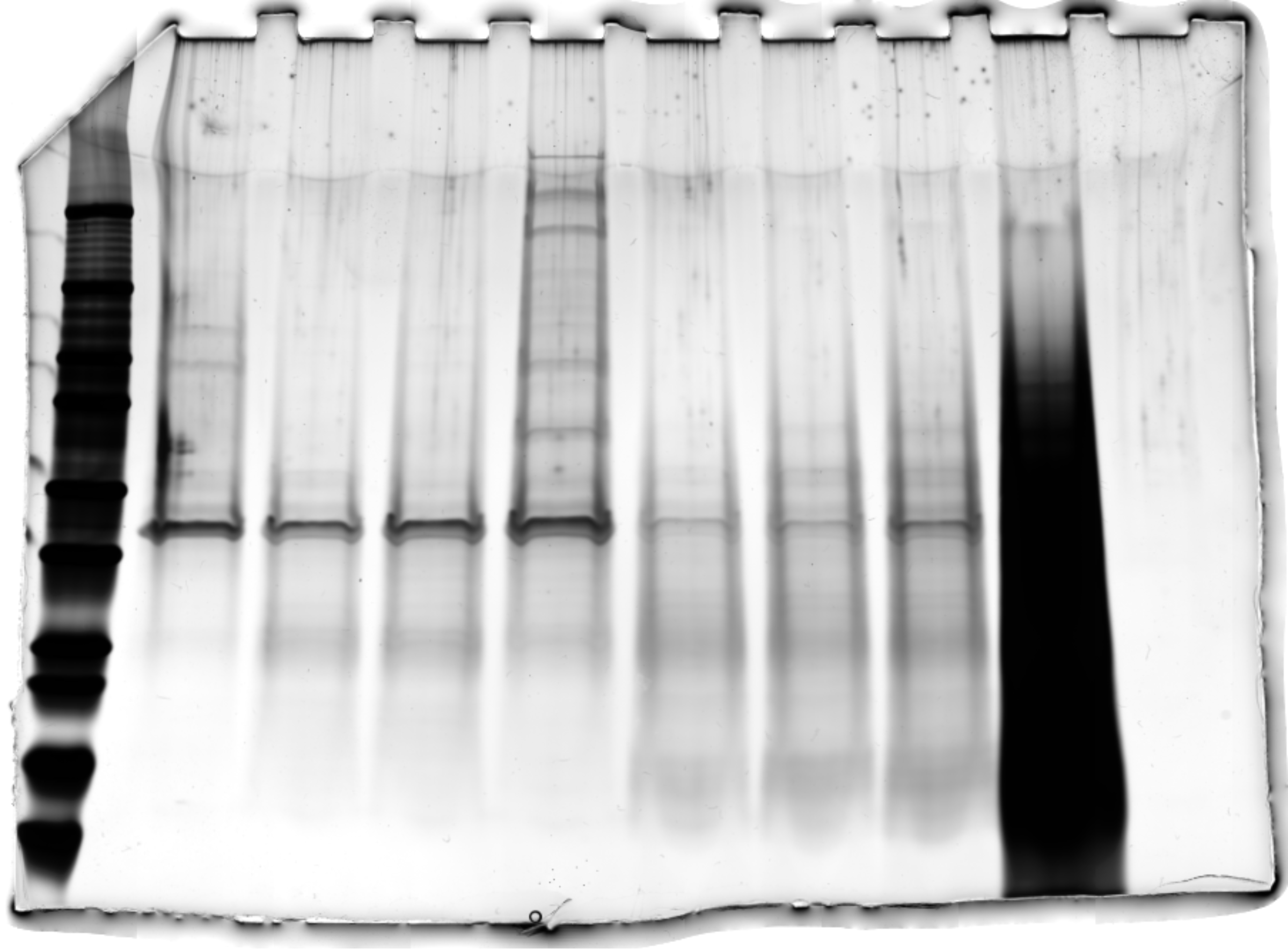
