## Supplementary material for "Impairment of lipid homeostasis causes lysosomal accumulation of endogenous protein aggregates through ESCRT disruption": Figure 3 - figure supplement 2-source data 1

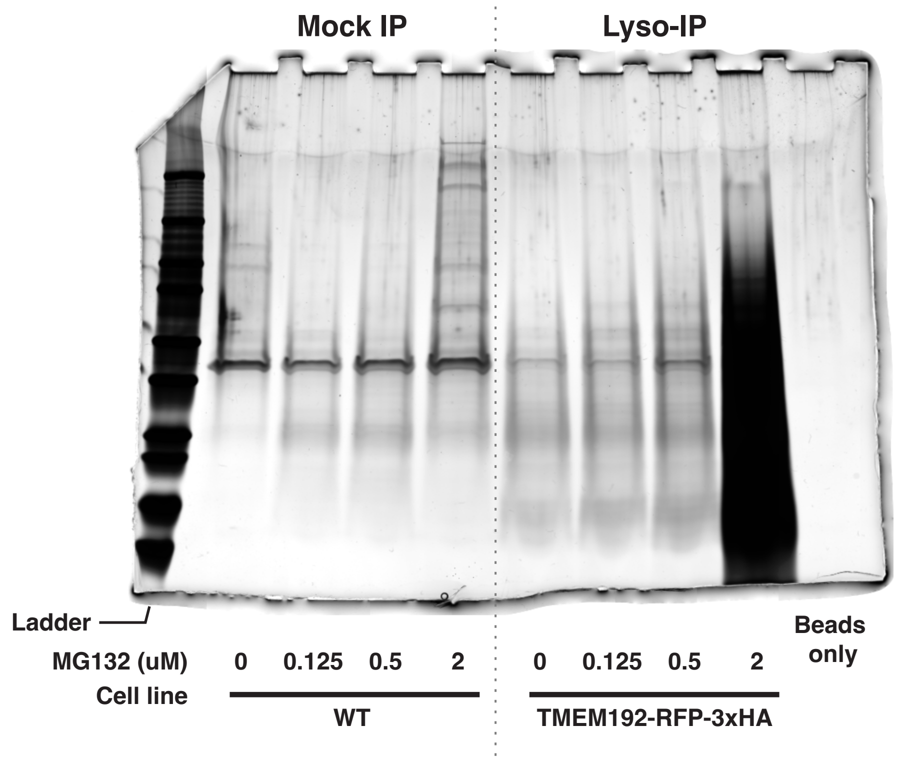


Ladder = BioRad Precision Plus Dual Color Protein Standards
