## Supplementary material for "Impairment of lipid homeostasis causes lysosomal accumulation of endogenous protein aggregates through ESCRT disruption": Figure 6-figure supplement 1-source data 1

**Blue epifluorescence signal of Me4BoVS labeling before cell lysis**

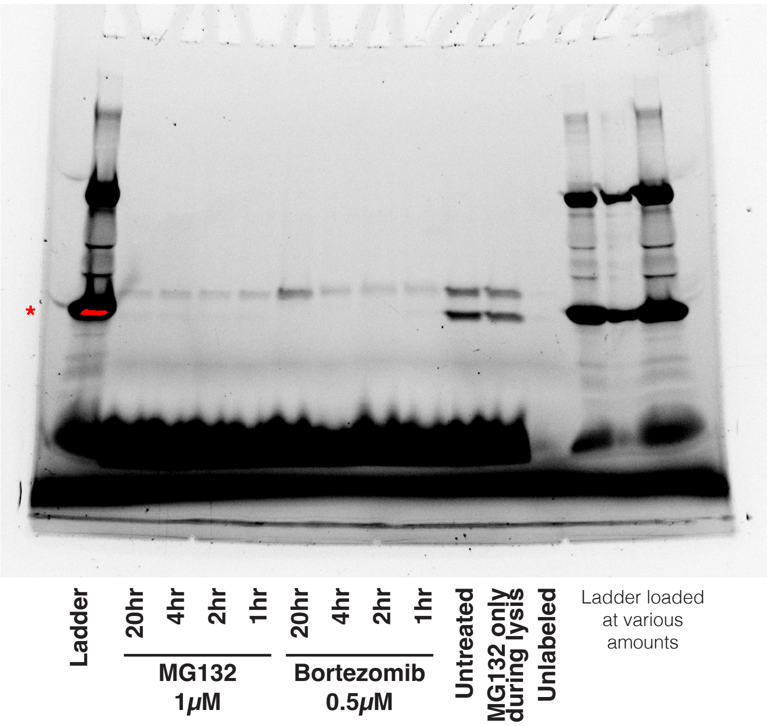


Ladder = BioRad Precision Plus Dual Color Protein Standards

* indicates signal saturation
